# A new model of Notch signaling: Control of Notch receptor cis-inhibition via Notch ligand dimers

**DOI:** 10.1101/2022.05.09.491117

**Authors:** Daipeng Chen, Zary Forghany, Xinxin Liu, Haijiang Wang, Roeland M.H. Merks, David A. Baker

**Affiliations:** School of Mathematics and Statistics, Xi’an Jiaotong University, Xi’an, China; Mathematical Institute, Leiden University, Leiden, The Netherlands; Leiden University Medical Center (LUMC), Department of Cell & Chemical Biology, Leiden, The Netherlands; Department of General Surgery, The First Affiliated Hospital, Xi’an Jiaotong University, Xi’an, China; Institute of Biology Leiden, Leiden University, Leiden, The Netherlands

## Abstract

All tissue development and replenishment relies upon the breaking of symmetries leading to the morphological and operational differentiation of progenitor cells into more specialized cells. One of the main engines driving this process is the Notch signal transduction pathway, a ubiquitous signalling system found in the vast majority of metazoan cell types characterized to date. Broadly speaking, Notch receptor activity is governed by a balance between two processes: 1) intercellular Notch transactivation triggered via interactions between receptors and ligands expressed in neighbouring cells; 2) intracellular cis inhibition caused by ligands binding to receptors within the same cell. Additionally, recent reports have also unveiled evidence of cis activation. Whilst context-dependent Notch receptor clustering has been hypothesized, to date, Notch signalling has been assumed to involve an interplay between receptor and ligand monomers. In this study, we demonstrate biochemically, through a mutational analysis of DLL4, both *in vitro* and in tissue culture cells, that Notch ligands can efficiently self-associate. We found that the membrane proximal EGF-like repeat of DLL4 was necessary and sufficient to promote oligomerization/dimerization. Mechanistically, our experimental evidence supports the view that DLL4 ligand dimerization is specifically required for cis-inhibition of Notch receptor activity. To further substantiate these findings, we have adapted and extended existing ordinary differential equation-based models of Notch signalling to take account of the ligand dimerization-dependent cis-inhibition reported here. Our new model faithfully recapitulates our experimental data and improves predictions based upon published data. Collectively, our work favours a model in which net output following Notch receptor/ligand binding results from ligand monomer-driven Notch receptor transactivation (and cis activation) counterposed by ligand dimer-mediated cis-inhibition.

**Author summary:** The growth and maintenance of tissues is a fundamental characteristic of metazoan life, controlled by a highly conserved core of cell signal transduction networks. One such pathway, the Notch signalling system, plays a unique role in these phenomena by orchestrating the generation of the phenotypic and genetic asymmetries which underlie tissue development and remodeling. At the molecular level, it achieves this via two specific types of receptor/ligand interaction: intercellular binding of receptors and ligands expressed in neighbouring cells, which triggers receptor activation (transactivation); intracellular receptor/ligand binding within the same cell which blocks receptor activation (cis inhibition). Together, these counterposed mechanisms determine the strength, the direction and the specificity of Notch signalling output. Whilst, the basic mechanisms of receptor transactivation have been delineated in some detail, the precise nature of cis inhibition has remained enigmatic. Through a combination of experimental approaches and computational modelling, in this study, we present a new model of Notch signalling in which ligand monomers promote Notch receptor transactivation, whereas cis inhibition is induced optimally via ligand dimers. This is the first model to include a concrete molecular distinction, in terms of ligand configuration, between the main branches of Notch signalling. Our model faithfully recapitulates both our presented experimental results as well as the recently published work of others, and provides a novel perspective for understanding Notch-regulated biological processes such as embryo development and angiogenesis.

## Introduction

The ubiquitous Notch pathway is an ancient, highly conserved signalling system whose early appearance in evolution coincided with the emergence of multicellularity (1,2). It was the first cell receptor signal transduction pathway to be discovered, more than a century ago, and decades of research since then have established that it is a central regulator of cell fate (1,2) that underpins normal embryo development and tissue homeostasis (3–8). Moreover, corruption of this network has been implicated in numerous pathologies including neurovascular diseases (CADASIL), multisystem disorders (ALAGILLE syndrome) (9) as well as the majority of solid tumours (10–12). Whilst invertebrates such as *Drosophila* possess a single Notch receptor family member controlled by two cognate ligands, in vertebrates, the Notch pathway is composed of up to four distinct receptor types (Notch1-4) and five different Type 1 transmembrane ligands: Jagged (JAG)1, JAG2, Delta-Like (DLL)1, DLL3, and DLL4 (13,14). Operationally, the canonical Notch signaling pathway is relatively well characterized. It is activated in juxtacrine manner through a *trans* interaction between single pass receptors expressed at the surface of one cell and ligands expressed by neighboring cells resulting in structural changes effected by biomechanical strain/pulling forces, which expose specific enzyme cleavage sites (15,16). Ultimately, a cascade of proteolytic events terminates in the γ-secretase-mediated cleavage of the Notch intracellular domain (17,18), which translocates to the nucleus whereupon it regulates expression of Notch target genes (19,20). In addition to transactivation, Notch is subject to another major regulatory mechanism termed cis-inhibition by which ligands block the activity of receptors expressed in the same cell (21,22). Collectively, these two counterposed processes (transactivation and cis-inhibition) are critical for determining the strength, the duration, the directionality and the specificity of Notch signalling.

In recent years, alongside cell, biochemical and genetic analyses, powerful mathematical approaches coupled to *in silico* modelling have become an important element of the toolkit needed to decipher the molecular details of Notch signalling and to understand the biological consequences of these processes (23–30). Collier et al. first proposed a mathematical description of lateral inhibition, an evolutionary conserved intercellular signalling mechanism that underlies symmetry breaking in tissues, in which Notch receptor (trans)activation in one cell via ligands expressed by neighbouring cells establishes the differential developmental cell fates necessary for patterning (23). Whereas this model can recapitulate essential features of transactivation, it was not until the work of Sprinzak et al., which has served as a common starting point for subsequent refinements, that cis inhibition and transactivation were integrated into a single model (22,25). Latterly, Elowitz and co-workers have developed a new model which takes account of the recently reported phenomenon of cis-activation (31). Whilst these technical and conceptual advancements are beginning to unravel the deeper complexities of Notch signalling, arguably a significant impediment to obtaining a more complete picture of this vital pathway is the relative paucity of the architectural/molecular details of cis and trans receptor/ligand complexes. Quantitative measurements of Notch/ligand binding have been performed, and structural studies have sought to identify specific binding interfaces (16,32), however, these analyses have relied upon investigating isolated receptor and ligand domains owing to the currently unsurmounted technical difficulties associated with purifying, and structurally and biophysically characterizing full length proteins. One consequence of this, in the absence of available evidence, is that it has been generally assumed that cis and trans receptor/ligand interactions are essentially monomeric. There are, however, sound reasons to suppose that the true picture may be more complicated. Both receptors and ligands harbour multiple EGF-like repeats, which are known to mediate protein-protein interactions (33). Related to that, we here show biochemically that Notch ligands can efficiently self-associate. This begged the question: what are the potential molecular and biological consequences of Notch ligand oligomerization? Through a combination of experimental approaches and mathematical modelling, we propose a novel view of Notch signalling in which ligand monomer-driven receptor transactivation (and cis-activation) is counterbalanced by ligand dimer-mediated cis inhibition.

## Results

### Notch ligands form dimers/oligomers

To date, it has been assumed that Notch ligands function as monomers. To formally explore this at the biochemical level, we first expressed epitope-tagged ligands in tissue culture cells and tested if the ligands could homo-oligomerize. Fig 1A shows that four different Notch ligands could efficiently self-associate. To further dissect the molecular basis of these interactions, we performed a detailed analysis of the DLL4 ligand. Fig 1B shows, in tissue culture cells, that the DLL4 extracellular domain is necessary and sufficient for homo-oligomerization and that plasma membrane anchorage is not required for this interaction. The DLL4 intracellular domain was found to be dispensable for DLL4-DLL4 binding (Fig 1B). We further demonstrate that whilst cis homo-oligomerization is very efficient (DLL4 molecules expressed in the same cell), trans oligomerization was not observed under the same conditions, though it cannot be ruled out that trans interactions were beyond the detection limit of the experiment (Fig 1C). Collectively, these data show that Notch ligands can forms oligomers both in tissue culture cells and also *in vitro*.

**Fig 1.**
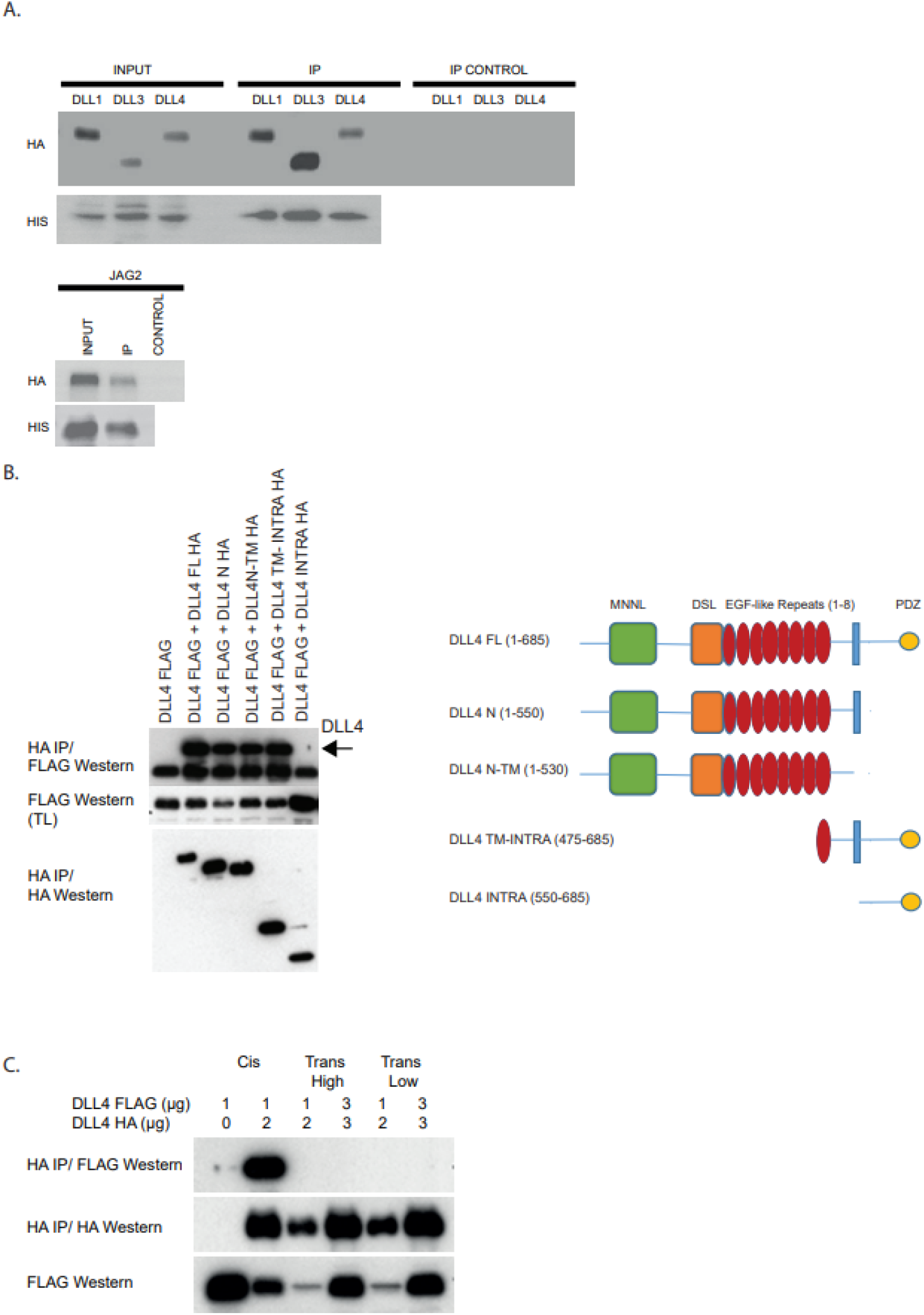
Notch ligand homo-oligomerization. (A) HIS epitope-tagged Notch ligands were purified from tissue-culture cells and incubated with the indicated HA-epitope tagged proteins produced by *in vitro* translation. Ligand-ligand interactions were determined by Western blotting using the shown antibodies. (B) Left panel: The indicated constructs were transfected into tissue-culture cells. Complexes were resolved by immunoprecipitation and visualized with the shown antibodies. Right panel: schematic representation of the constructs used in the study. (C) The indicated constructs were transfected into tissue-culture cells in one of three ways: cis-ligands were co-expressed in the same cells sparsely plated to exclude trans interactions; trans (high)-differently tagged ligands were expressed individually in cells, which were subsequently mixed in confluent cell monolayers to enable trans interactions; trans (low)-as for trans (high) but cells were plated at low cell density. Complexes were resolved by immunoprecipitation and visualized using the shown antibodies.

### The membrane proximal DLL4 (EGF-like repeat 8) and the MNNL domain bind to DLL4 *in vitro*

To identify the domains responsible for DLL4 oligomerization, we performed a comprehensive mapping analysis using purified proteins. By these means, we found that EGF-like repeat 8 but not EGF-like repeats 1-7, either as part of the DLL4 extracellular domain (see Fig 2A and 2B) or singularly (Fig 2C), binds to DLL4, consistent with the idea that this domain might underpin DLL4-DLL4 binding (see Fig 2). These experiments also highlighted the MNNL domain as binding efficiently to DLL4 (Fig 2B).

**Fig 2.**
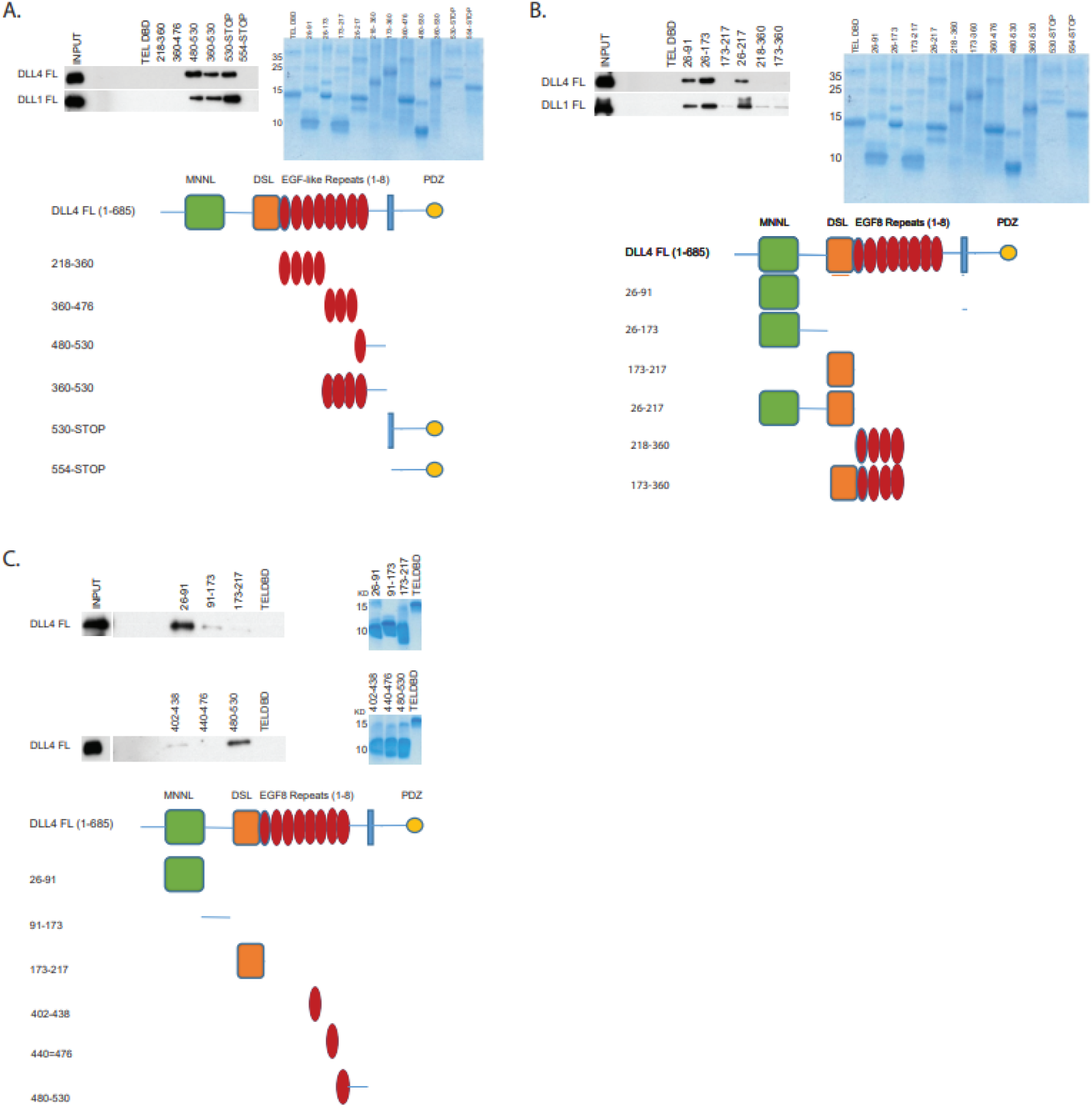
Biochemical mapping of DLL4 dimerization motifs. (A-C) The indicated HIS-epitope tagged proteins (a schematic representation of constructs is shown) were purified from *E. coli* (representative Coomassie-stained gels of protein preparations are shown) and incubated with full length DLL4 ligand manufactured by *in vitro* translation. Ligand-ligand Interactions were determined by Western blotting.

### EGF-like repeat 8 mediates DLL4-DLL4 binding but is dispensable for DLL4 binding to the Notch receptor

To test the requirement of the EGF-like repeat 8 and the MNNL domain for DLL4 oligomerization, we expressed epitope-tagged wild type and mutant DLL4 ligands in tissue culture cells. Whereas the MNNL domain was found to be dispensable for DLL4-DLL4 binding under these conditions, loss of the EGF-like repeat 8 abrogated binding (Fig 3A). Moreover, deletion of EGF-like repeat 7 or EGF-like repeat 6 did not detectably inhibit ligand-ligand binding suggesting that the EGF-like repeat 8 encodes a specific DLL4 oligomerization motif (Fig 3B) and that deletion of the EGF-like repeat did not non-specifically corrupt ligand-ligand binding. To determine the impact of deleting the EGF-like repeat 8 on ligand-receptor binding, we co-expressed wild type or mutant DLL4 ligands with Notch 2. Fig 3C shows that DLL4 mutants lacking either EGF-like repeat 8, EGF-like repeat 7, EGF-like repeat 6 or the MNNL domain, associated with Notch2 as efficiently as wild type DLL4. Together, these findings support the view that the DLL4 EGF-like repeat 8 specifically mediates DLL4-DLL4 binding but is not required for Notch receptor-DLL4 binding. Since mutant DLL4 harbouring a deletion of EGF-like repeat 8 presumably exists primarily as a monomer, these results suggest that ligand oligomerization is not a general pre-requisite for Notch receptor binding.

**Fig 3.**
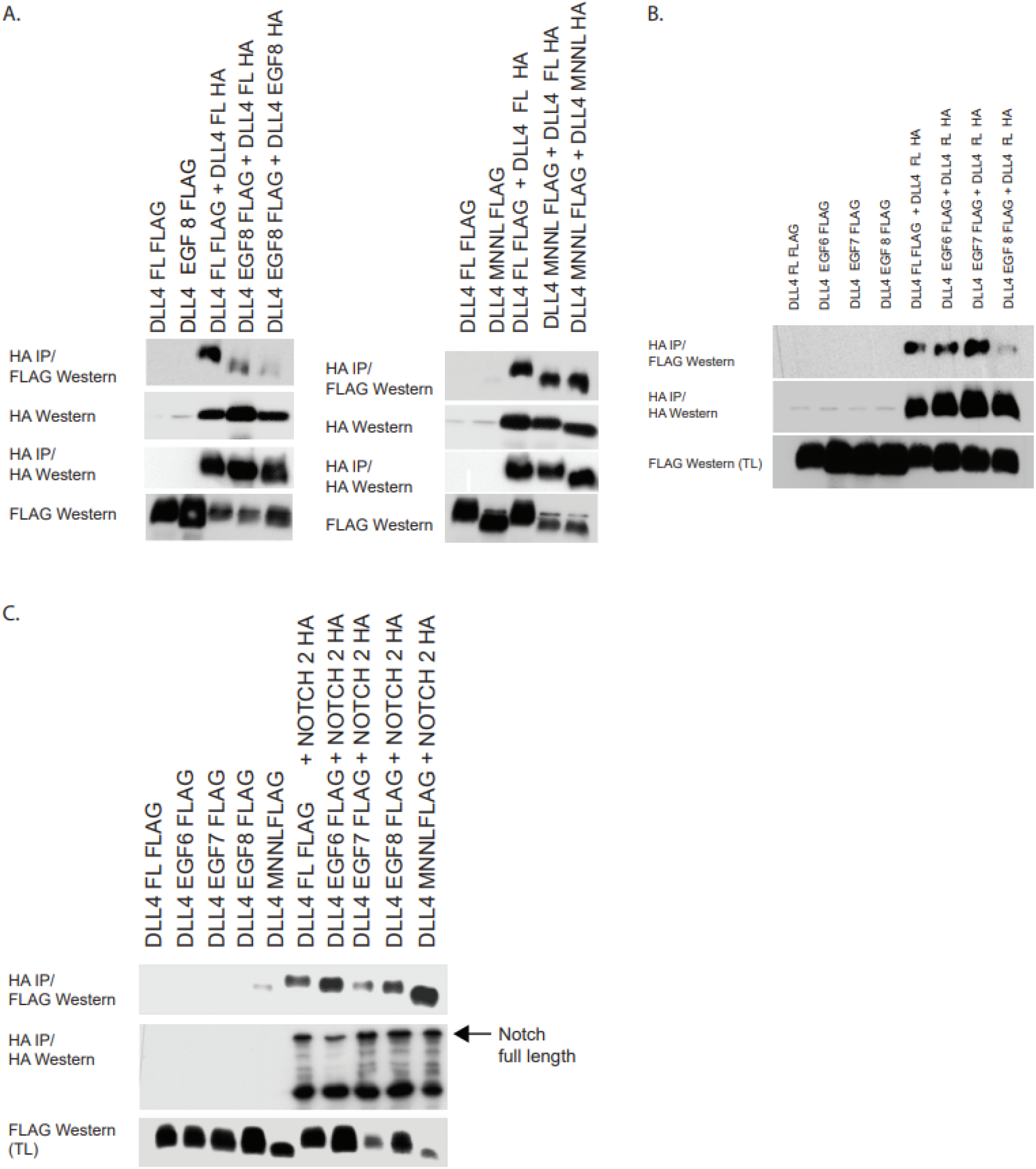
(A-C) The indicated constructs were transfected into tissue-culture cells. DLL4EGF6, EFG7, EGF8, MNNL each have deletions of the named domain. Complexes were resolved by immunoprecipitation and visualized by Western blotting with the highlighted antibodies.

### DLL4 oligomerization is required for cis-inhibition of the Notch receptor

To elucidate the mechanistic consequences of ligand oligomerization, we performed luciferase reporter assays to quantitatively measure Notch receptor activity. When expressed in the same cell as Notch2 receptors, wild type DLL4 or mutant DLL4 ligands lacking either EGF-like repeat 7 or EGF-like repeat 6, all of which can form oligomers, inhibited the activity of Notch2 approximately 5-fold. By contrast, DLL4 ligands lacking EGF-like repeat 8, which thus act as monomers, failed to inhibit Notch2 activity when co-expressed in the same cell (Fig 4A). When Notch2 and wild type DLL4 (or mutant DLL4) were expressed in neighbouring cells, we found that Notch receptor transactivation was unaffected by deletions of EGF-like repeat 8, EGF-like repeat 7 or EGF-like repeat 6 (see Fig 4A). Fig 4B shows that deletion of EGF-like repeat 8 did not result in any overt change in the sub-cellular location of DLL4. Overall, these results favour a model in which cis-inhibition, but not transactivation, of Notch signalling specifically depends upon DLL4 ligand oligomerization.

**Fig 4.**
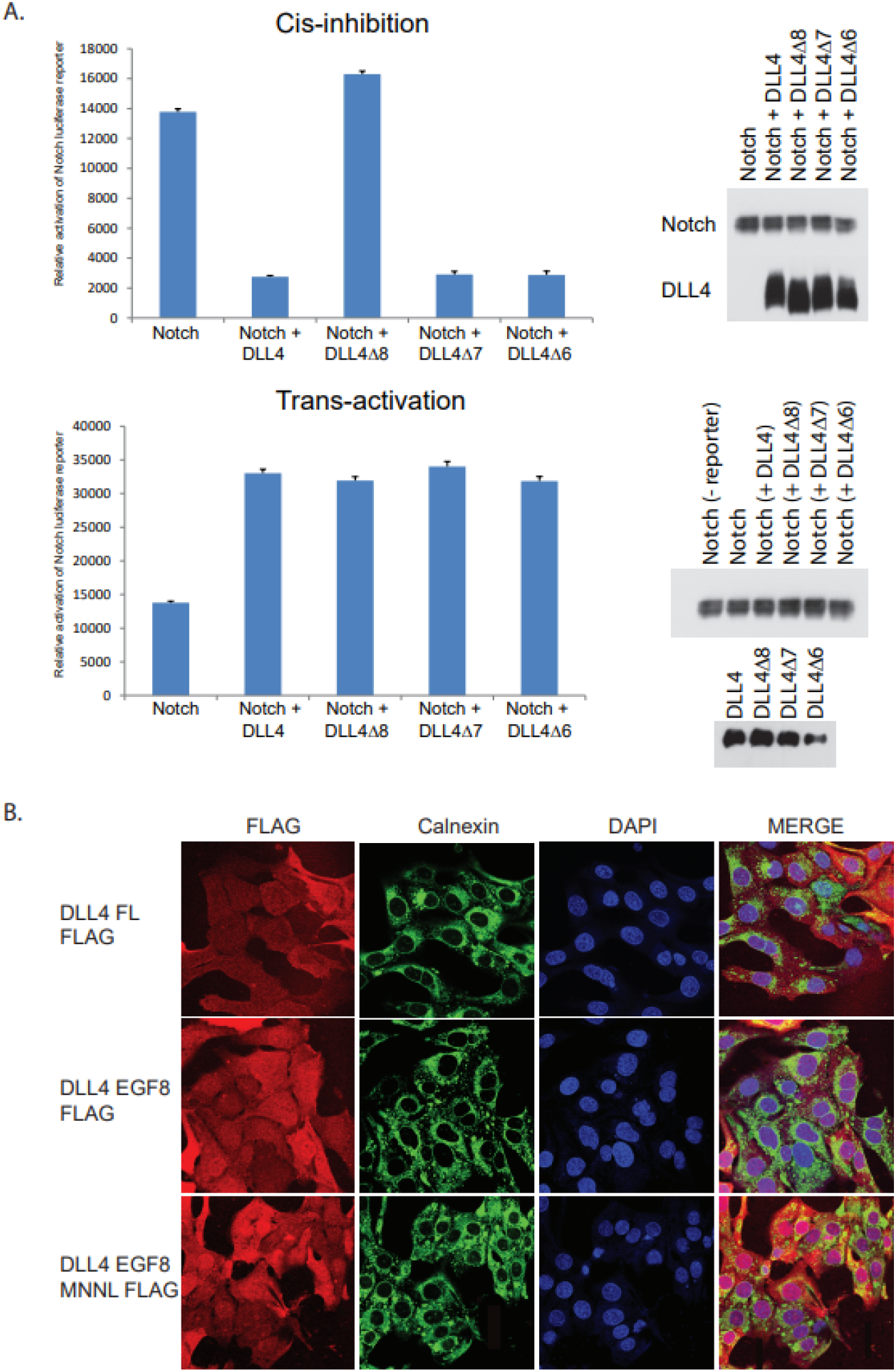
(A) Luciferase reporter assays. Upper graph: U2OS cells were co-transfected with a Notch activity luciferase reporter together with the indicated Notch2 and ligand constructs. Lower panel: Cells transfected with Notch2 and a Notch activity luciferase reporter were separately co-cultured with cells expressing the indicated DLL4 constructs. For each analysis, reporter activity was normalized using Renilla luciferase. Levels of ectopically expressed proteins were determined by Western blotting of cell lysates. Experiments were performed in triplicate. (B) Immunofluorescence showing the sub-cellular distribution of wild type and mutant DLL4 ligands.

### A novel mathematical model demonstrating ligand oligomer-dependent cis-inhibition of Notch signalling

To further explore the potential biological implications of our biochemical findings presented above, we have adapted the mutual inactivation model recently proposed by Sprinzak et al. (22) to include ligand oligomerization and cis-activation (see S1 Text). In contrast to their model, in which cis-inhibition is driven via ligand monomers, in our model cis-inhibition is driven via ligand oligomers. The corresponding mathematical model (1) includes a new compartment *L_n_*, indicating the amount of ligand oligomers composed of *n* ligand monomers, *L*,

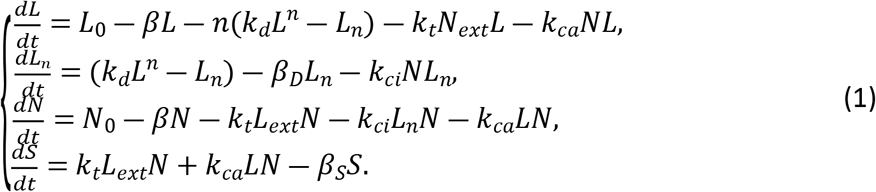

where *L*_0_ and *N*_0_ denote the net expression levels of Notch ligand and Notch receptor (*N*), respectively. The proteins in Model (1) are assumed to be degraded at a constant rate given by *β, β_D_* and *β_s_*, which describe the degradation rates of protein monomer, ligand oligomer and the Notch intracellular domain (*S*), respectively. A comprehensive description of all parameters is detailed in Table 1 and Supplementary Information (S1 Text).

**TABLE 1.** DESCRIPTION AND BASELINE VALUES OF PARAMETERS USED IN SIMULATIONS.

| PARAMETER | Description | Values | Units | Source |
| --- | --- | --- | --- | --- |
| $L_0$ | Baseline production rate of Notch ligands | 200 | molec * hour<sup>-1</sup> | Estimated from (27) |
| $N_0$ | Baseline production rate of Notch receptors | 200 | molec * hour<sup>-1</sup> | Estimated from (27) |
| $k_d$ | Baseline oligomerization rate of ligand monomers | $1 * 10^{-3}$ | molec<sup>-(n-1)</sup> * | Assumed |
| $k_t$ | Trans-activation rate | $5 * 10^{-5}$ | molec<sup>-1</sup> * hou | (26,27,28) |
| $k_{ci}$ | Cis-inhibition rate | $6 * 10^{-4}$ | molec<sup>-1</sup> * hou | (26,27,28) |
| $k_{ca}$ | Cis-activation rate | $5 * 10^{-6}$ | molec<sup>-1</sup> * hou | Estimated from (31) |
| $\beta$ | Degradation rate of protein monomers | 0.1 | hour<sup>-1</sup> | (26,27,28) |
| $\beta_D$ | Degradation rate of Notch ligand oligomers | 0.01 | hour<sup>-1</sup> | Assumed |
| $\beta_S$ | Degradation rate of free Notch Intracellular Domain | 0.5 | hour<sup>-1</sup> | (26,27,28) |

In Model (1), Notch receptors can be trans-activated via ligands (*L_ext_*) expressed in neighboring cells or cis-inhibited by ligands expressed in the same cell. The model makes the following initial assumptions: transactivation of Notch is mediated by ligand monomers; both monomers and oligomers can bind to Notch in cis resulting in either cis-inhibition (via ligand oligomers) or cis-activation (via ligand monomers); ligand oligomerization is a reversible process. The data presented in Fig 4 have shown that ligand monomers are sufficient to maximally stimulate Notch transactivation and ligand oligomers are necessary to mediate Notch receptor cis-inhibition. To explore other potential roles of ligand oligomers in Notch signalling, we compared Model (1) with a number of alternative models. Among the model variants, we found that Model (1) best matches the experimental data. For simplicity, in the following analyses, we initially focus on Notch ligand dimerization (*n* = 2 in Model 1). A detailed comparison of the effect of ligand dimers (denoted as *L*\*) versus higher order ligand oligomers (n > 2) on Notch activity is given later (see Fig 8), which showed that ligand dimer is the configuration which best fits the experimental data.

### Exploring the role of ligand monomers and ligand dimers in trans-activation and cis-inhibition

To test the validity and robustness of our model, as a first step, we performed simulations to establish if the model could recapitulate the experimental findings described in Fig 4, that is, could the model demonstrate a requirement for ligand oligomer-dependent cis-inhibition. Fig 5A schematically depicts two alternative general cases: that ligand dimers specifically and exclusively mediate cis-inhibition (see case 1 in Fig 5A); that ligand dimers mediate cis-inhibition and trans-activation (see case 2 in Fig 5A). These cases are described mathematically by,

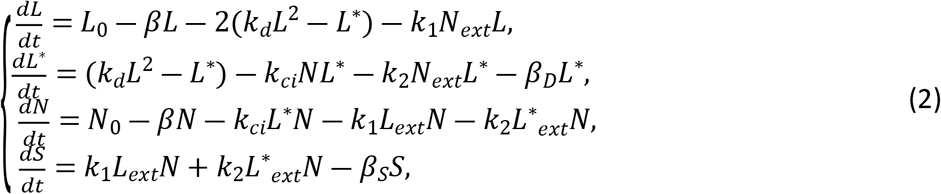

with *k*_1_ = *k_t_* and *k*_2_ = 0 for case 1; *k*_1_ = *k_t_* and *k*_2_ = *k_t_* for case 2. Here we do not consider cis-activation because cis-activation is much weaker than trans-activation (31).

**Fig 5.**
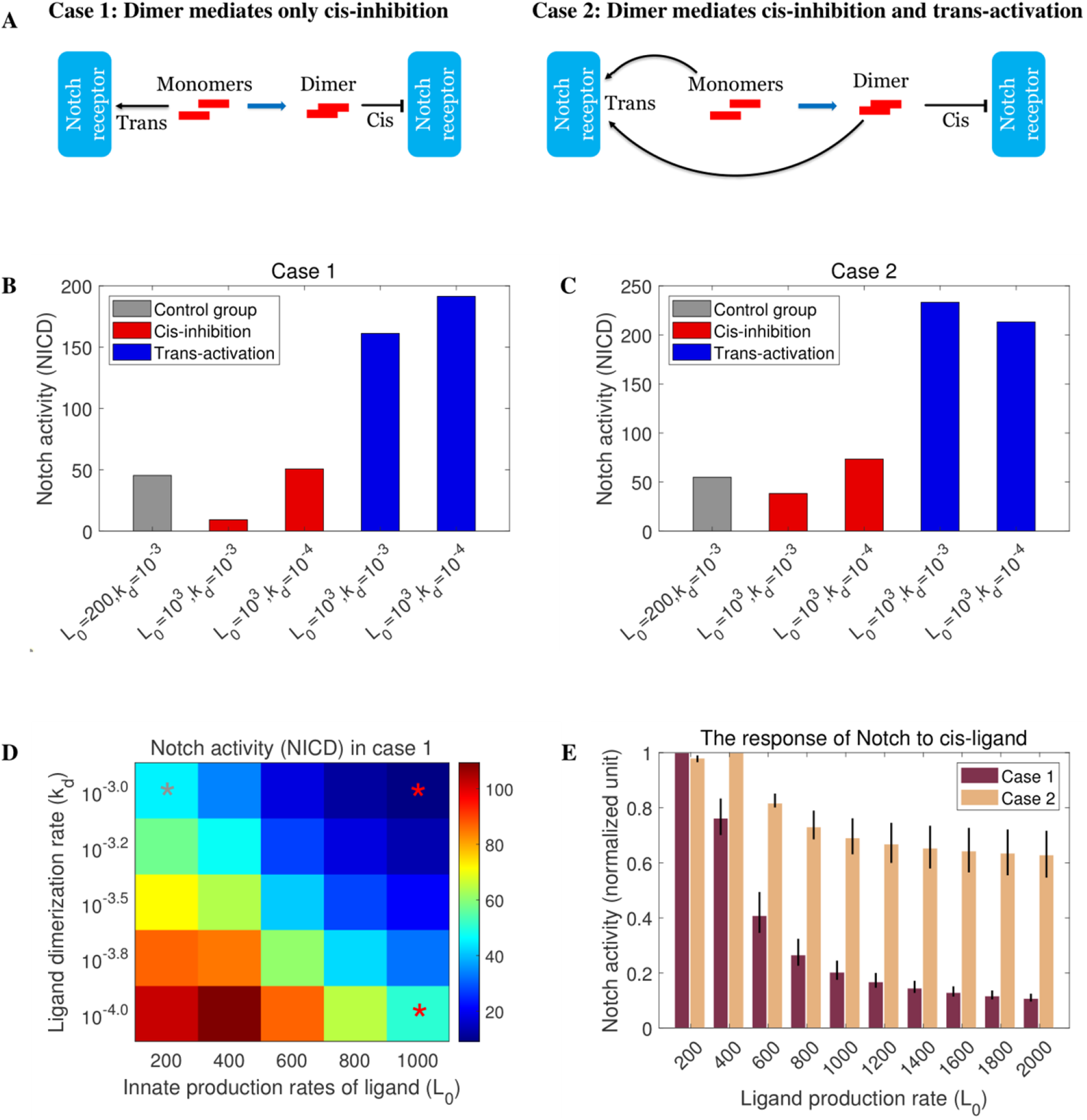
The role of Notch ligand monomer and dimer in trans-activation and cis-inhibition. (A) The potential rules governing protein interactions in Notch signaling. Case 1: ligand dimer only mediates cis-inhibition; case 2: ligand dimer mediates both cis-inhibition and trans-activation. (B-C) In silico replication of cell-based luciferase experiments shown in Fig 4A. Ligand production rate (*L*_0_) and dimerization rate (*k_d_*) are varied in the simulations. The grey bar represents a control group with baseline parameters. The red groups with high ligand production rate (*L*_0_) in two adjacent cells recapitulates cis-inhibition. The blue groups with high ligand production rate (*L*_0_) in one of two adjacent cells recapitulates trans-activation. (D) The heatmap of Notch activity across a broad range of Notch ligand production rates and dimerization rates. Three conditions used in panels B and C are marked with, *. (E) Relative Notch activity in response to increasing Notch ligand production rates for the two cases shown in A. Notch activity is normalized against the maximum possible activity. The range of ligand dimerization rates (*k_d_*) is between 10^−2^ and 10^−4^ /(molec * hour).

We considered two adjacent cells and fixed the ligand production rate, *L*_0_, and the ligand dimerization rate, *k_d_*, based upon previously published estimates (see Table 1 and S1 Text). Fig 5B and 5C show that case 1 (Fig 5B) most closely recapitulated the experimental data (Fig 4A). In case 1, high levels of ligand leads to a strong reduction in Notch activity via cis-inhibition, whereas high levels of cis-ligand in case 2 (Fig 5C) cannot inhibit Notch activity effectively because of competition with ligand dimers-mediated trans-activation, suggesting that ligand dimerization specifically promotes cis-inhibition. Two additional numerical analyses lend further support to this view. Firstly, Fig 5D shows that case 1 (see Fig 5A) holds true over a broad range of *L*_0_ and *k_d_* values. Secondly, Fig 5E shows that over a broad range of ligand dimerization rates (values were based upon published ligand-specific molecular interactions), case 1 but not case 2 reproduces ligand dimer-dependent cis-inhibition in a ligand concentration-dependent manner, as would be expected, that is, higher ligand expression levels elicit greater levels of cis-inhibition. In summary, multiple unbiased simulations, utilizing a broad range of physiologically relevant parameters, demonstrate that the mathematical model (1), describing ligand-oligomer-specific cis-inhibition of Notch signalling, faithfully recapitulates our experimental findings.

To test the general applicability of our model, we ran simulations of cis-inhibition and trans-activation of Notch signalling to test if it could reproduce previously published experimental data (22). To measure these phenomena experimentally, Sprinzak et al. deployed elegant cell-based reporter assays enabling quantification of both cis-inhibition and trans-activation of Notch signalling (22). We compared the accuracy of their mutual inactivation model (Model 5 in the Methods section) with our ligand dimerization model (case 1 in Model 2) using an experimental data set that demonstrated unambiguously cis-inhibition of Notch signalling (shown as an inset in Fig 6A). Our ligand dimerization model (case 1 in Model 2) also successfully reproduces the reported experimental findings, indeed it appears to more closely match the observed pattern of ligand decay (Fig 6B; Fig SI 1C in S1 Text). We extended these studies by investigating Notch trans-activation. We derived the relationship between Notch activity (quantified by the production rate of fluorescence) and trans-ligand level based on Model (1),

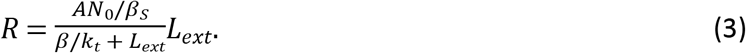

where *A* is a scale parameter that converts protein level to fluorescence (see Methods for details). Fig 6C and 6D show the parameterization of this relationship and identify a curve of best fit which approximate the observed experimental Notch trans-activation results.

**Fig 6.**
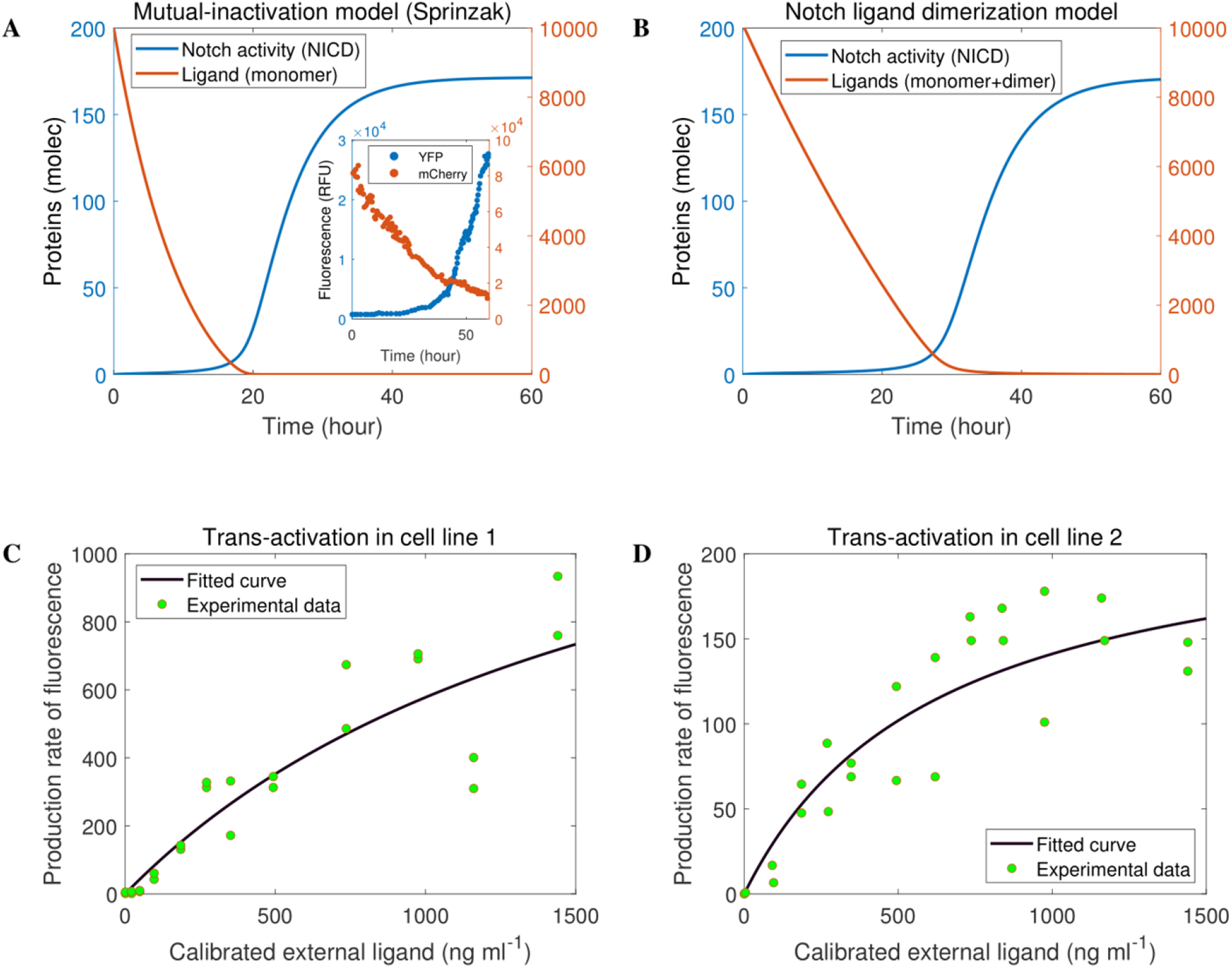
Testing the general applicability of our model on the dynamics of cis-inhibition and trans-activation in Notch signaling. (A) Experimental data and simulations of cis-inhibition using the mutual-inactivation model (22). (B) Simulations to cis-inhibition using our Notch ligand dimerization model. (C-D) Eq 3 was fitted to the observed experimental Notch trans-activation results, respectively. The fitting parameters are two combined parameters *AN*_0_/*β_s_* and *β/k_t_* in Eq (3).

### Modelling the role of ligand monomers and ligand dimers in cis-activation and cis-inhibition

Elowitz and co-workers recently reported cis-activation as a novel, previously overlooked Notch signalling mechanism (31). To investigate the potential role of ligand dimerization in this process, we proposed three possible scenarios (cases 1 to 3 in Fig 7A) based on the general Model (1), given mathematically by,

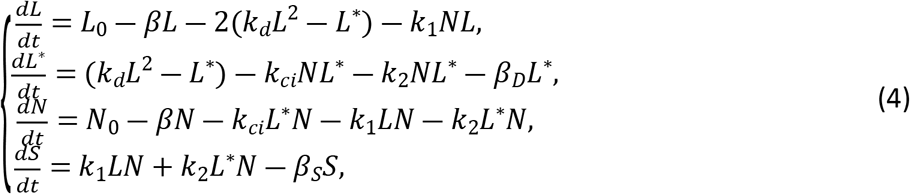

with *k*_1_ = *k_ca_* and *k*_2_ = 0 for case 1; *k*_1_ = 0 and *k*_2_ = *k_ca_* for case 2; *k*_1_ = *k_ca_* and *k*_2_ = *k_ca_* for case 3. Using cell-based assays, Nandagopal et al. (31) observed a non-monotonic response of Notch to cis ligand levels (Fig 7B). To explain this response, they proposed a number of possible cases. The model which most closely reproduced their experimental observations suggests that cis-activation occurs through formation of a ligand-receptor complex, *C*, and that binding of a further ligand to the complex leads to cis-inhibition (their model 2c), given mathematically by Model (6) in the Methods section. By running simulations using reference parameter values (Table 1) based upon their experimental results (Fig 7B), in Fig 7C-E we demonstrated that among several proposed ligand dimerization models, the experimental data is best and only explained by Model (4) with case 1 (compare Fig 7B and 7C-7E), which assumes that ligand dimers mediate cis-inhibition whilst ligand monomers mediate cis-activation (see case 1 in Fig 7A). The mechanism proposed by Nandagopal et al. (31) also reproduces the experimental data (compare Fig 7B and 7F), but does not include the explicit ligand oligomerization step that our data suggest. Notably, our ligand dimerization model generates narrower peaks using the same reference parameter values, which is closer to experimental data (compare Fig 7B, 7C and 7F). Interestingly, the rate of ligand dimerization might modulate both the width and the amplitude of the Notch cis-activation peak, as well as the ligand concentration at which optimal cis-activation is reached (Fig 7G). In summary, our simulations suggest that cis-activation is mediated by ligand monomers (Fig 7C), but the non-monotonic response of Notch to cis ligand levels is dependent on ligand dimer-mediated cis-inhibition (Fig 7G).

**Fig 7.**
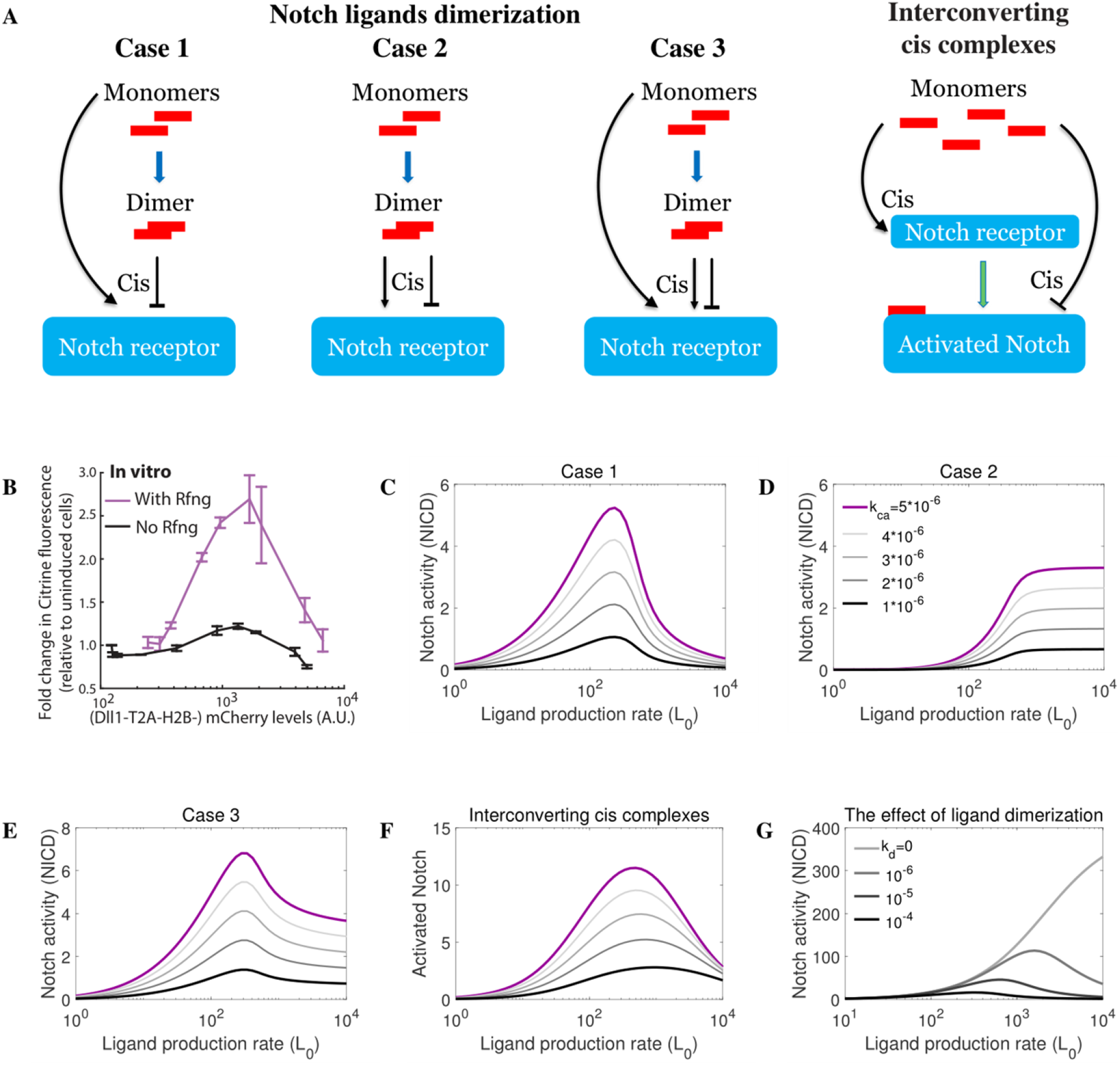
The role of Notch ligand monomers and dimers in cis-activation. (A) Schematic representation of different potential rules governing receptor/ligand interactions in Notch signaling. Case 1: monomer mediates cis-activation and dimer mediates cis-inhibition; case 2: dimer mediates cis-activation and cis-inhibition; case 3: monomer mediates cis-activation, dimer mediates both cis-activation and cis-inhibition; a model proposed by Nandagopal et al. (31) in which monomers mediate cis-activation and cis-inhibition in two consecutive steps. (B) Published in vitro cis-activation experiments (31). The response of Notch to cis ligand level is non-monotonic. (C-F) Simulations of cis-activation for each of the cases using our Model 4 (C-E) or the model proposed by Nandagopal et al. (31) (F). Different cis-activation rates are tested. (G) The role of Notch ligand dimerization in cis-activation. Notch ligand dimer does not directly mediate cis-activation, but ligand dimerization is necessary to explain the experimental observations.

**Fig 8.**
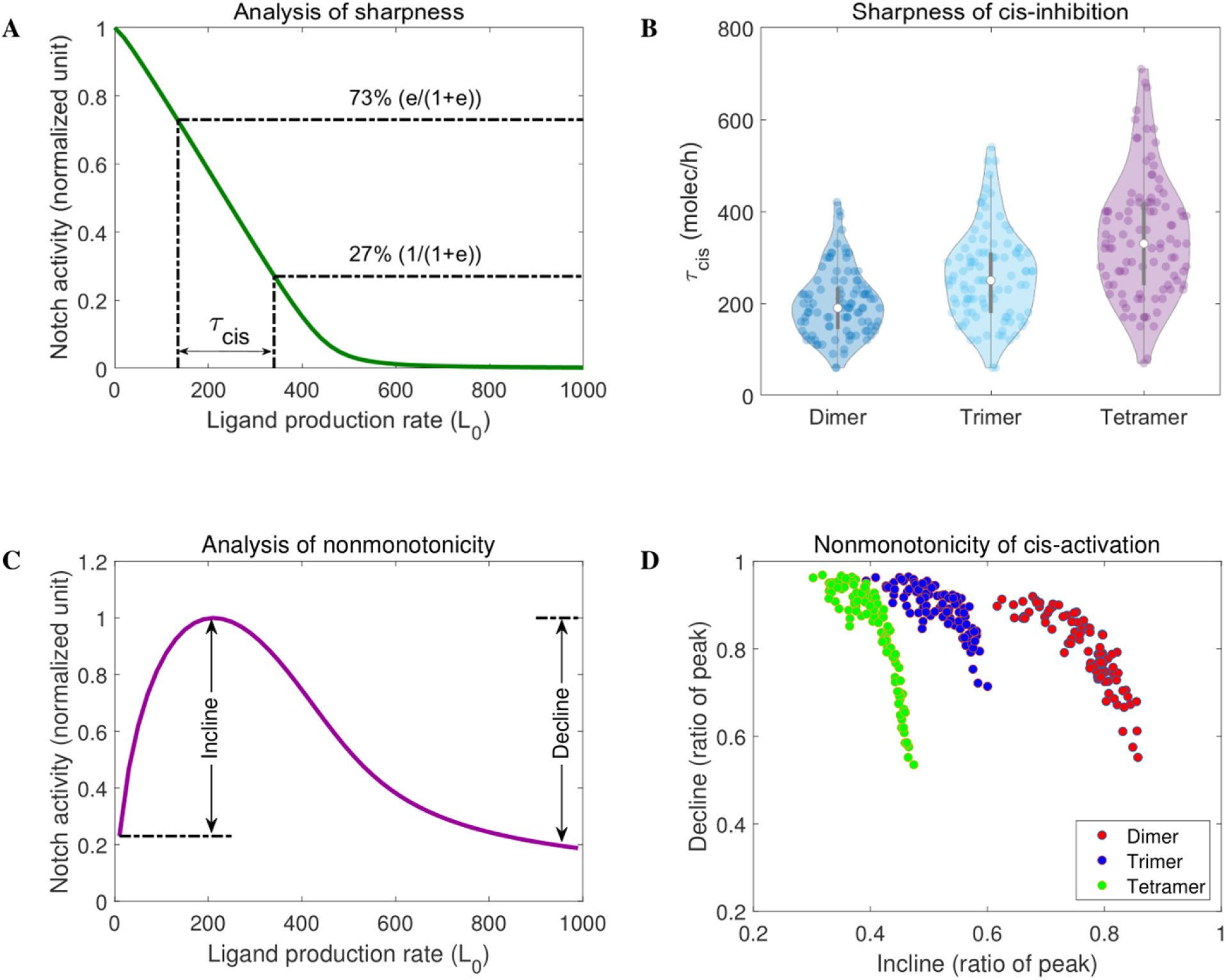
The effect of ligand oligomer size on cis-inhibition and cis-activation. (A) The response of Notch to cis-ligand defines a quantity, (*τ_cis_*), used to measure the sharpness of cis-inhibition. (B) The effect of ligand oligomer size on the sharpness of cis-inhibition showing dimer-dependent cis-inhibition is optimally sharp and robust. (C) Definition of two quantities (incline and decline) used to measure the nonmonotonicity of cis-activation. (D) The effect of ligand oligomer size on cis-activation nonmonotonicity. Dimer-dependent cis-inhibition generates a robust pattern of nonmonotonic cis-activation.

### The effect of ligand oligomer size on cis-inhibition and cis-activation

It is currently not possible biochemically to quantitatively distinguish between ligand homo-oligomerization and homo-dimerization. In Fig 8, we consider mathematically the ligand oligomer configuration which best describes the observed experimental data. Using our general Model (1) in combination with reported experimental data (22), we first derived a term, *τ_cis_*, to describe the overall sharpness, that is, the rate and strength of cis-inhibition (Fig 8A). Next, we determined this value over of broad range of parameters by running simulations of cis-inhibition mediated by ligand oligomers of different size (Fig 8B). These analyses clearly demonstrate that ligand-dimer-dependent cis inhibition, in contrast to cis-inhibition directed by ligand oligomers of greater size, most closely resembles the results of published experimental studies (22), in which the estimated sharpness of cis-inhibition is equivalent to the Hill function of cis-ligand with Hill coefficient 12. We extended these findings by considering the effect of ligand oligomer size on the pattern of cis-activation. We defined two quantities, incline and decline, to describe the increasing range and decreasing range of Notch activity in response to cis-ligand (Fig 8C). Next, we determined these values over a broad range of parameters by running simulations of cis-activation controlled by ligand oligomers of different size (see Fig 8D). Again, the analyses clearly showed that dimers, but not other oligomers, are best to explain the observed non-monotonic pattern of cis-activation in response to cis-ligand. This most closely resembles the results of published experimental studies (31).

## Discussion

In this study, we have taken a combined experimental and mathematical modelling approach to establish biochemically that vertebrate Notch ligands can self-associate, and to dissect the potential role of this phenomenon in Notch signal transduction. Several lines of evidence are presented, centred chiefly on the Notch ligand, DLL4, which support the view that net signalling output from the Notch pathway is the product of ligand monomer-mediated transactivation (and cis-activation) counterposed by ligand oligomer-mediated cis inhibition of Notch receptor activity. Our new model is the first to propose a concrete molecular distinction, in terms of ligand configuration, between the two principal branches of Notch signalling, specifically that whereas ligand monomers promote Notch receptor transactivation, they are insufficient to account for cis inhibition which is induced optimally via ligand dimers. Our model exhibits two features that are consistent with our molecular findings (see Figs 1-3). Firstly, cis inhibition is driven by ligand dimers, which can associate independently of receptor binding prior to receptor/ligand complex formation, that is, cis inhibition is not (strictly) a stepwise process of monomer to dimer transitions assembled on preformed receptor/ligand complexes at the cell membrane. Secondly, our modelling results do not depend on cis inhibition occurring at a particular subcellular compartment, such as the cell membrane. Rather, cis inhibition can also occur cytoplasmically (Fig SI 1 in S1 Text), in line with our observations that ligands lacking a transmembrane domain can efficiently self-associate and execute Notch receptor cis inhibition (see Fig 1 & Fig4; ZF & DAB unpublished data). These data are in agreement with earlier studies demonstrating cytoplasmic receptor/ligand binding (38) and raise the possibility that cytoplasmic ligand dimer-dependent cis inhibition of Notch activity could serve to block ‘mis-firing’ of the receptor prior to its expression at the plasma membrane. Related to this, whilst it is clear that ligand dimers are necessary to drive cis inhibition, it not clear whether both ligand dimers and monomers are capable of stimulating receptor transactivation. To address this mathematically, we considered two cases: dimers promote cis inhibition/monomers promote transactivation; dimers promote cis inhibition/monomers and dimers can promote transactivation (Fig 5A). Using physiologically relevant parameters our simulations strongly favour the former case (see Fig 5). In further support of this idea is our finding that ligands harbouring mutations that abrogate self-association, stimulated Notch transactivation as efficiently as wild type ligands (see Fig 4). The general applicability of our model was demonstrated by the fact that it can also recapitulate the results of previously published work (see Fig 6).

Nandagopal et al. recently unveiled a previously overlooked dimension of Notch signalling, termed cis activation, which results from monomeric interactions between receptors and ligands expressed at the plasma membrane of the same cell (31). In agreement with this, although we cannot formally demonstrate that our dimerization-defective mutant can elicit wild type levels of cis activation, we consistently observed enhanced reporter activity under cis conditions in the presence of DLL4 ligand monomer but not DLL4 ligand dimer, which instead strongly inhibited Notch receptor activity (see Fig 4), albeit in the absence of Radical Fringe which reportedly enhances receptor/ligand interactions and could have correspondingly enhanced these effects. Furthermore, our model could faithfully recapitulate the published experimental data (see Fig 7). Intriguingly, their mathematical model postulates that cis activation becomes cis inhibition, in a ligand concentration-dependent fashion, following binding of additional ligand monomers to ligand monomers complexed with Notch receptors at the cell membrane (presumably prior to receptor-bound monomer promoting cis activation). In light of this, and given the results of our own simulations, our model suggests that ligand dimerization may ensure a correct balance between cis inhibition and cis activation/trans activation (see Fig 7G).

To date, a major gap in Notch signalling knowledge is the precise nature of ligand, receptor and receptor/ligand complexes at a detailed molecular/structural level encompassing protein-protein interactions and the precise role of post-translation modifications such as glycosylation (39). In particular, it is unknown if trans receptor/ligand interactions differ conformationally from cis receptor/ligand interactions, and whether or not this could underlie or at least contribute to their distinct effects on receptor activity. Whilst such architectural details are currently lacking, a notable prediction of our mathematical modelling is that ligand dimers, rather than larger ligand oligomers, should be optimal for controlling cis inhibition of Notch receptor activity (see Fig 8). Additionally, Notch ligand oligomerization could have major implications for our understanding of the dynamics of the Notch pathway because regulation of dimer formation/dimer disassembly, might represent an extra point of Notch signalling strength control.

There are four mammalian Notch receptors and five mammalian Notch ligands, and the manifold potential receptor-ligand combinations will give rise to the different signalling outputs necessary for tissue patterning. It is established that different ligands can elicit unique cell fates. By example, JAG1 and DLL4 exert opposing effects on angiogenesis (40). Moreover, recent work has established that ligands can stimulate either discrete pulses of Notch activity (in the case of DLL1) or a sustained period of signalling (in the case of DLL4) yielding distinct gene expression outcomes (24). In this context, two other potential facets of Notch signalling merit consideration. Whilst here we have uncovered a mechanistic role for ligand oligomerization, there is growing evidence that Notch receptors can also oligomerize/dimerize though the precise molecular consequences of this remains elusive (41,42). Moreover, since all Notch ligands share a common overall architecture, it could be of interest to investigate if there is a biological role for ligand hetero-oligomerization. Interestingly, simple biochemical experiments revealed that ligands can indeed hetero-oligomerize (see Fig SI 2 in S1 text). Such investigations coupled to refined mathematical models will help to fully disentangle this core signalling pathway.

In summary, we have delineated a previously unreported requirement for ligand dimerization in the cis inhibition of Notch receptor activity. This new mechanism could help determine the strength, the direction, the specificity and the nature of the output of the Notch signalling pathway. A novel mathematical model has been developed which successfully captures this process at the molecular level. It will be of interest to test if the same model can provide important insights into Notch-controlled physiological processes such as sprouting angiogenesis (43–45), and, given the global interest in the role of Notch signalling in human pathogenesis (46,47), to determine how these ideas will impact the design of novel therapeutic approaches to diseases.

## Methods

### Mathematical modelling

#### A formula for trans-activation

For engineering cells in which the expression of Notch ligand was inhibited (*L*_0_ = 0) (22), model (1) implies that the level of both Notch ligand monomer and oligomers is equal to zero. Therefore, the non-negative equilibrium (0,0,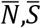) of system (1) satisfies

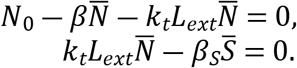

Based on these equations, the stable Notch activity is given by

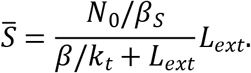

In experiments, a fluorescent protein related to Notch-activity is often used to quantify the Notch activity since the time series of fluorescence intensity *F*(*t*) can be obtained by tracking living cells. The fluorescent protein is relative stable, so we assume that the fluorescence (*F*(*t*)) of cells is proportional to cumulative level of NICD (denoted as *S*). Consequently, 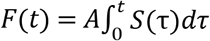, where *A* is a transfer constant from protein level to fluorescence. Therefore, the production rate of fluorescence is *R*(*t*) = *F*’(*t*) = *AS*(*t*), which is the formular (Eq 3). Based on the data on trans-activation obtained by Sprinzak et al. (22), we estimate the parameters in the formula from 100 iterations of fitting 8 time points, chosen randomly with replacement, out of a total 12 measured time points.

#### Mutual inactivation model (22)

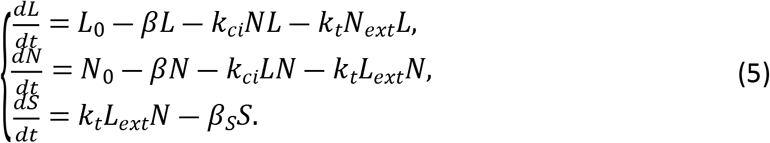

#### Interconverting cis complexes model (31)

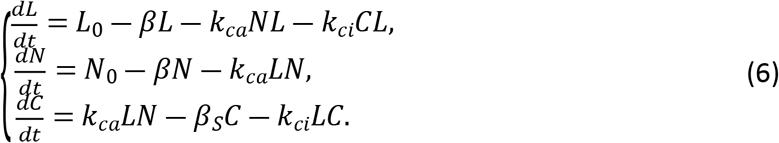

Where *C* is the activated ligand-Notch complex which could be inhibited by ligand monomer in their model (31).

#### Simulation details

Numerical simulation, nonlinear curve-fitting and numerical analysis were performed on MATLAB R2021a. Runge–Kutta methods were used in the numerical solutions of all models. Nonlinear least-squares method was used to fit Notch trans-activation by the Eq (3). Latin hypercube sampling was used to generate the parameter sets.

**Fig 5 and Fig SI 1A-B:** To mimic the interactions in experiments, we consider two adjacent cells. In this context, the external ligand (*L_ext_*) and receptor (*N_ext_*) in models (1) and (2) come from neighboring cell. That is to say, here we run a system with eight equations. Considering cis-inhibition, we changed ligand production rate (*L*_0_) and ligand dimerization rate (*k_d_*) in two cells simultaneously. To test trans-activation, we only changed ligand production rate (*L*_0_) and ligand dimerization rate (*k_d_*) in one of adjacent cells.

**Fig 6 and Fig S1 1C:** Here we focus on an individual cell exposed to fixed external ligand (22). Similar to experiments, the production rate of Notch ligand is zero. To simulate the observed cis-inhibition, we set *k_ci_* = 6 * 10^−3^ in Fig 6A-B and Fig SI 1C, and set high initial number of ligands (10000 molecules, 0 molecules, 0 molecules, 0 molecules) in Fig 6A, (2500 molecules, 7500 molecules, 0 molecules, 0 molecules) in Fig 6B and (2500 molecules, 7500 molecules, 0 molecules, 0 molecules, 0 molecules, 0 molecules) in Fig S1C.

**Fig 7 and Fig SI 1D:** Cis-activation appears to be weaker than trans-activation. Consequently, we tested different cis-activation rates (*k_ca_*) smaller than the reference of trans-activation rate.

**Fig 8:** We generated 100 parameter sets using Latin hypercube sampling in the hypercube whose center is baseline parameters. In each parameter dimension, we consider a range increasing and decreasing by 50%. We ran our model repeatedly with different parameter sets.

### Experimental methods

#### Cell culture, biochemistry and molecular biology

Human embryonic kidney 293T cells and U2OS osteosarcoma cells were cultured in DMEM (Gibco) supplemented with 10% fetal bovine serum (Gibco). Cell lines were typed using short tandem repeat analysis of the DNA and all cell lines were checked for mycoplasma with the MycoAlert kit (Lonza). Transfections, lentivirus production and cell infections, Western blotting and co-immunoprecipitations have been described previously (34,35). All lysis buffers contained a cocktail of protease inhibitors (phenylmethylsulfonyl fluoride, trypsin inhibitor, pepstatin A, leupeptin, aprotinin).

#### Recombinant protein production/ *in vitro* protein:protein interaction

Domains for recombinant protein production were cloned into the pET 28a vector in-frame to an N-terminal 6x HIS epitope. His epitope-tagged proteins were manufactured in *Escherichia coli* BL21(DE3). Following sonication (Misonix Sonicator 3000) in 3 mls ice-cold buffer / 50 ml bacterial culture (150 mM NaCl, 2.7 mM KCl, Na_2_HPO_4_, KH_2_PO_4_, 20 mM imidazole, 10 mM *β*-mercaptoethanol), proteins were purified onto 50 ul of Nickel-agarose beads (Qiagen) by 3 hours rolling at 4C. Beads were washed in 10 x 1 ml of the same buffer. Protein yields were determined by Bradford assay (Bio-Rad) and relative protein integrity and purity was determined by SDS-PAGE and Colloidal Blue staining (Invitrogen). Purified recombinant protein was incubated with 10 ul nickel beads in 1 ml of buffer for 2 hours at 4°C with *in vitro* translated DLL4 proteins made using the TNT-coupled reticulocyte *in vitro* translation system (Promega). Beads were washed x10 with 1 ml of buffer. Proteins were separated by SDS-PAGE and associated proteins were detected by Western blot.

#### Plasmid construction

Unless otherwise stated, all cDNAs were fused in-frame with a Flag or an HA epitope tag and were cloned into the pLV lentiviral vector and pCS2 expression plasmid. Mutants were generated by site-directed mutagenesis using Phusion High-Fidelity DNA polymerase (Thermo fisher). All constructs were verified by Sanger sequencing (Macrogen).

#### Immunofluorescence

Immunostaining was performed as previously described (36,37) using Alexa Fluor 488 goat anti-mouse secondary antibodies (Thermo Fisher scientific). Imaging was performed with a Leica SP8 confocal microscope.

#### Luciferase reporter

Cells ectopically expressing the relevant receptor/ligand combinations were seeded in triplicate in 12-well plates and transfected with 2ug of a Notch luciferase reporter harbouring 10x RBPJ consensus binding sites, and Renilla luciferase control plasmid. Cells were lysed 36 hours post-transfection, and luciferase activity was measured using a luciferase assay substrate (Promega). Luciferase activity was normalized by measuring Renilla luciferase activity (Promega). Experiments were performed three times.

#### Antibodies and drugs

Antibodies were obtained from the following sources: FLAG mouse M2 monoclonal (Sigma); anti-HA.11 mouse monoclonal (Covance); anti-HA rabbit polyclonal (Abcam); anti-FLAG rabbit (Sigma); anti-γ-tubulin (Sigma); anti-GFP (GeneTex); anti-His (Sigma). Anti-Calnexin (AbCam).

## Supporting information

**S1 Text. Supplementary information of this paper.** The supplementary document provides modeling details, parameters description, three supplementary figures and MATLAB code for the main text.

## Acknowledgements

We thank members of the Departments of Cell & Chemical Biology and the Leiden Institute of Biology for helpful discussions, technical advice and support, in particular Professor Peter ten Dijke and Jacob Visscher.

